# Cast away on Mindoro island: lack of space limits population growth of the endangered tamaraw (*Bubalus mindorensis*)

**DOI:** 10.1101/2020.05.07.051037

**Authors:** C. Bonenfant, A. Rutschmann, J. Burton, R. Boyles, F. García, A. Tilker, E. Schütz

## Abstract

Endangered species, despite often living at low population densities, may undergo unexpected density-dependent feedbacks in the case of successful recovery or marked reduction in range. Because density-dependence dynamics can increase risk of extinction, these effects can hamper conservation efforts. In this study, we analyse the dynamics of the largest population of the tamaraw (*Bubalus mindorensis*), a critically endangered ungulate species endemic to Mindoro island, Philippines. The population is located within a < 3,000 ha area in Mounts Iglit-Baco Natural Park, with limited expansion possibilities. We took advantage of a 21 year time series of tamaraw counts to estimate annual population growth rate and possible density-dependence, accounting for sampling errors in the counts. The tamaraw population has been increasing at an average rate of +5% per year, as would be expected given its protected status by law. Population growth showed strong spatial structuring, with a population growth close to +10% in the core area of protection, and a reduction of abundance of −5% at the periphery of its range, inside the protected area. This range constriction is concerning because our best population dynamics model suggests significant negative density-dependence (Bayes factor = 0.9). The contraction of tamaraw range is likely caused by anthropogenic pressures forcing the species to live at relatively high densities in the core zone where protection is most effective, creating source-sink dynamics. Our study highlights the fact that, despite the continuous population growth over the last two decades, the long-term viability of the Mounts Iglit-Baco Natural Park tamaraw population remains uncertain.

## INTRODUCTION

Assessing population abundance is an important component of conservation planning because of the strong link between population size and likelihood of extinction (Boyce 1992). Accordingly, low population abundance is one of the primary criteria used for the evaluation of conservation status (Mace *et al*. 2008; Neel *et al*. 2012). Indeed, the effects of small population size on species persistence has been identified as one of the two central paradigms of conservation biology (Caughley 1994). Research has shown that the population dynamics of small populations differ in several ways from what is reported at higher abundance (Caughley 1994; Mugabo et al. 2013). When population size is small, demographic stochasticity – the random death of a few individuals – is more influential on the population dynamics and its viability than in populations that are larger in size (Shaffer 1981; Lande 1993). Similarly, when mating partners are too few to meet and reproduction fails, demographic stochasticity generates the so-called Allee effect (Courchamp et al. 1999), characterized by a decrease of population growth at low population abundance (*i.e*. demographic component, see Stephens *et al*. (1999)). Yet demographic stochasticity and the Allee effect are not the only considerations for the conservation of small populations; other density-dependent effects may also put these vulnerable populations at risk.

Classically, negative density-dependence describes situations where population growth rate is altered by increasing animal density (Nicholson 1933), with crowding driving the fate and viability of populations at risk of extinction (Beissinger & Westphal 1998; Henle *et al*. 2004). Negative density-dependent effects on population dynamics and demographic rates are often overlooked in conservation for a simple reason: they are *a priori* expected to occur at high population abundance, a counter-intuitive situation for threatened populations. Yet, even in small populations, increasing abundance can occur rapidly as the result of (i) a range contraction caused by habitat loss, (ii) the reintroduction of individuals for population reinforcement, (*iii*) the disruption of natural emigration due to habitat fragmentation, or (iv) following successful protection measures without the possibility for the population to expand geographically. In situations where there is low carrying capacity and limited connectivity for animals to seek less crowed areas, negative density-dependence may hinder some conservation goals. Achieving long-term population growth, or the removal of animals for reinforcement are, for instance, deemed to be difficult when negative density-dependence is at work.

There are multiple well-documented density-dependence effects on vital rates (Eberhardt 1977; 2002; Fowler 1987; Bonenfant *et al*. 2009), most of which are the result of behaviour and life history trait changes as population density approaches carrying capacity (Bonenfant *et al*. 2009; Choquenot 1991; Van der Wal et *al*. 2014). First, with increasing population density, individuals may die from starvation because of a reduction in the availability and accessibility of food resources by conspecifics. Such antagonistic interactions may generate high stress levels which, in turns, magnify the effect of other mortality sources (Peterson & Black 1988; Hone & Clutton-Brock 2007). For instance, at high population density, epizootics are more likely to emerge and to spread quickly in populations (Langwig *et al*. 2012). Second, higher densities may also force non-dominant animals to occupy low-quality territories, ultimately decreasing the average productivity and survival of the population (Krüger & Lindström 2001). Given its numerous biological consequences, early detection of density-dependence is important for the management of threatened species because its evidence from data would suggest that the population is approaching the carrying capacity of the environment, and starts to be limited by food resources (Eberhardt 2002).

The tamaraw (*Bubalus mindorensis*) is a large ungulate endemic to the island of Mindoro, Philippines (Heude 1888; Custodio *et al*. 1996). The species was originally found across the entire island of Mindoro, with a total estimate of 10 000 individuals in 1900 (Long *et al*. 2018). Since then, tamaraw populations suffered from land conversion for agriculture and logging, trophy hunting, disease outbreaks spread by domestic cattle (Maala 2001), and traditional hunting conducted by upland indigenous communities who share their living space with the species (Long *et al*. 2018). Despite being officially protected by law since 1954, as well as the creation of the sizeable Mounts Iglit-Baco Natural Park (MIBNP) protected area in 1970 to protect its key habitat (Maala 2001), tamaraw range has decreased substantially over the last decades. As a consequence, the tamaraw is now found in a few isolated populations scattered across Mindoro, with MIBNP holding the largest population (Matsubayashi *et al*. 2010; Wilson & Mittermeier 2011; Long *et al*. 2018), restricted to a single location <3 000 ha hereafter referred to as the core zone of monitoring (CZM).

Given its importance for the conservation of this iconic species, the MIBNP tamaraw population has been the cornerstone of past, present, and future conservation plans (Maala 2001; Long *et al*. 2018). In an attempt to reinforce tamaraw protection, wildlife managers came to an agreement with residing indigenous communities in 2016, declaring a 1 600 ha no-hunting zone area within the CZM. Unfortunately, one unforeseen consequence of the no-hunting area establishment is an increase in hunting pressure at its boundaries. Park rangers regularly report the setting of snare and spear traps in the border areas, and violation of the no-hunting zone agreement is not uncommon, with several deaths of tamaraws confirmed in the last years (Boyles pers. comm.). Furthermore, field observations by rangers suggest that tamaraw population density is growing within the no-hunting zone, which could increase the risks of crowding. In this context, it is possible that negative density-dependence combined with the source-sink dynamics will affect the population in the near future, undermining long-term conservation strategies.

Here, we took advantage of a 21 year time series of abundance data to provide a detailed analysis of the tamaraw population dynamics in the CZM of MIBNP. We predict (*i*) that because of the substantial conservation effort in the last decade, overall population growth rate (noted *r*) should be statistically >0. Yet (*ii*), because the no-hunting agreement area creates a spatial demarcation between risky and safe habitats, we expect local growth rates to differ in space, and predict a decrease of r in areas further from the main rangers’ base camps and patrolling routes. Finally (iii), given the intensive conservation efforts in the CZM, and relative safety of tamaraw in the CZM, we predict a negative association between r and tamaraw abundance within the no-hunting zone as a result of negative density dependence effects. To test the first two hypotheses, we computed the population growth rate of the tamaraw population at two spatial scales: over the entire CZM, and at specific locations used in annual tamaraw counts (hereafter referred to as vantage points). To test our third hypothesis of density-dependent demographic responses, we fitted and compared three population dynamics models: the baseline exponential model (no density-dependence), a Ricker model to test for a linear relationship between *r* and annual abundance, and a particular formulation of Gompertz model to test for a decrease of *r* through time.

## MATERIAL AND METHODS

### Study site

The Mounts Iglit-Baco Natural Park (hereafter noted MIBNP) is a 106 655 ha protected area of the Philippines in the south-central part of the island of Mindoro (N12°54’, E121°13’). The Mounts Iglit-Baco Natural Park harbors the largest population of tamaraw, found in a single 3 000 ha area located on the south-west edge of the protected area. This 3 000 ha area demarcates our study site, and is where most patrolling and observation activities are carried out (the so-called CZM). There are currently around 15 park rangers, which are divided into two or three teams to monitor the area on a regular basis. The teams patrol routes within the CZM, centered around the three permanent base camps. The CZM and no-hunting zone represent approx. 3 and 1.5% of MIBNP, respectively.

The CZM area is a rolling grassland plateau with an average elevation of 800 m a.s.l., dominated by cogon (*Imperata cylindrica*) and wild sugarcane (*Saccharum spontaneum*), and interspersed with numerous wooded creeks, secondary forest fragments, and steep hills. Introduced from Borneo to Mindoro in the early 60s (Pancho & Plucknett 1971), the invasive Siam weed (*Chromolaena odorata*) has been rapidly spreading in the grassland area and now covers an unknown fraction of the tamaraw habitat. The tropical climate is strongly seasonal with a rainy season from June to October (mean temperature 26.5°C; mean rainfall: 150 mm), and a hot dry season from November to May (mean temperature 29.8°C; mean rainfall: 5 mm). Apart from humans, the only potential predator to tamaraw is the reticulated python (*Malayopython reticulatus*) that is capable of preying upon calves or yearlings, though no such predation observations have been made so far.

### Tamaraw counts

Every year since 2000, the Tamaraw Conservation Program (TCP), a program of the Philippine Department of Environment and Natural Resources (DENR), has conducted tamaraw counts across the CZM, yielding an annual index of relative population abundance. The tamaraw counting area consists of 18 vantage points, covering nearly 2 200 ha within the CZM (Fig. 2). The tamaraw counting process is rather invasive for the MIBNP ecosystem because it involves the burning of grasslands a few weeks ahead of each count to increase visibility and to retain tamaraw to areas with nutrient-rich young grasses. For the past 21 years, between 1 200 and 1 500 ha of grasslands were intentionally burned each year for the purpose of monitoring tamaraw abundance.

**Figure 1.**
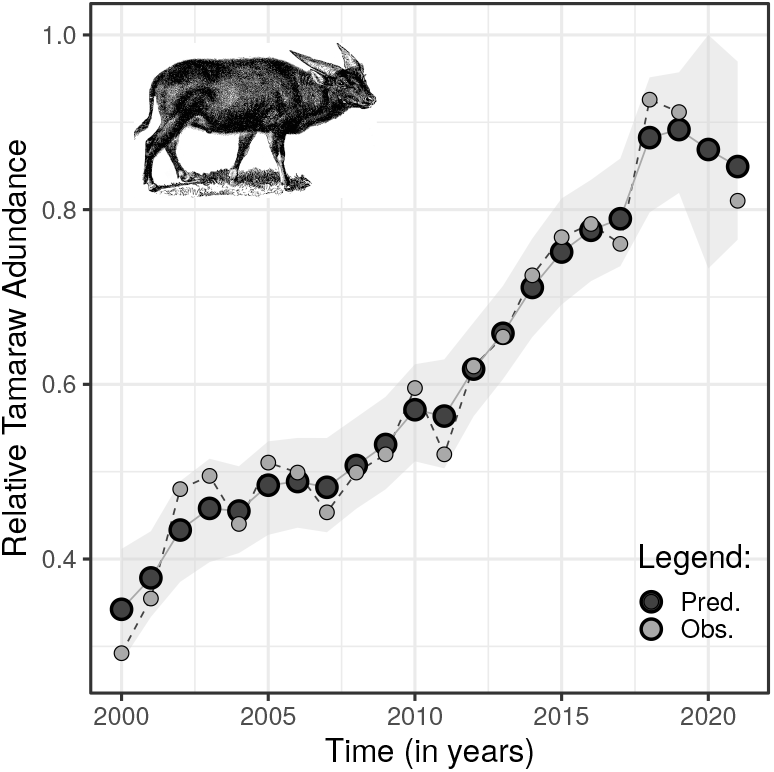
Time series of tamaraw (*Bubalus mindorensis*) abundance in Mounts Iglit-Baco Natural Park on Mindoro island, Philippines, from years 2000 to 2021 (with the exception of 2020 because of the COVID pandemic). Raw counts (light symbols) and counts corrected for sampling variance (dark symbols) are presented with the associated 95% credible intervals. Over the 21 years of monitoring, tamaraw abundance increased at a rate of 5% per year on average.

**Figure 2.**
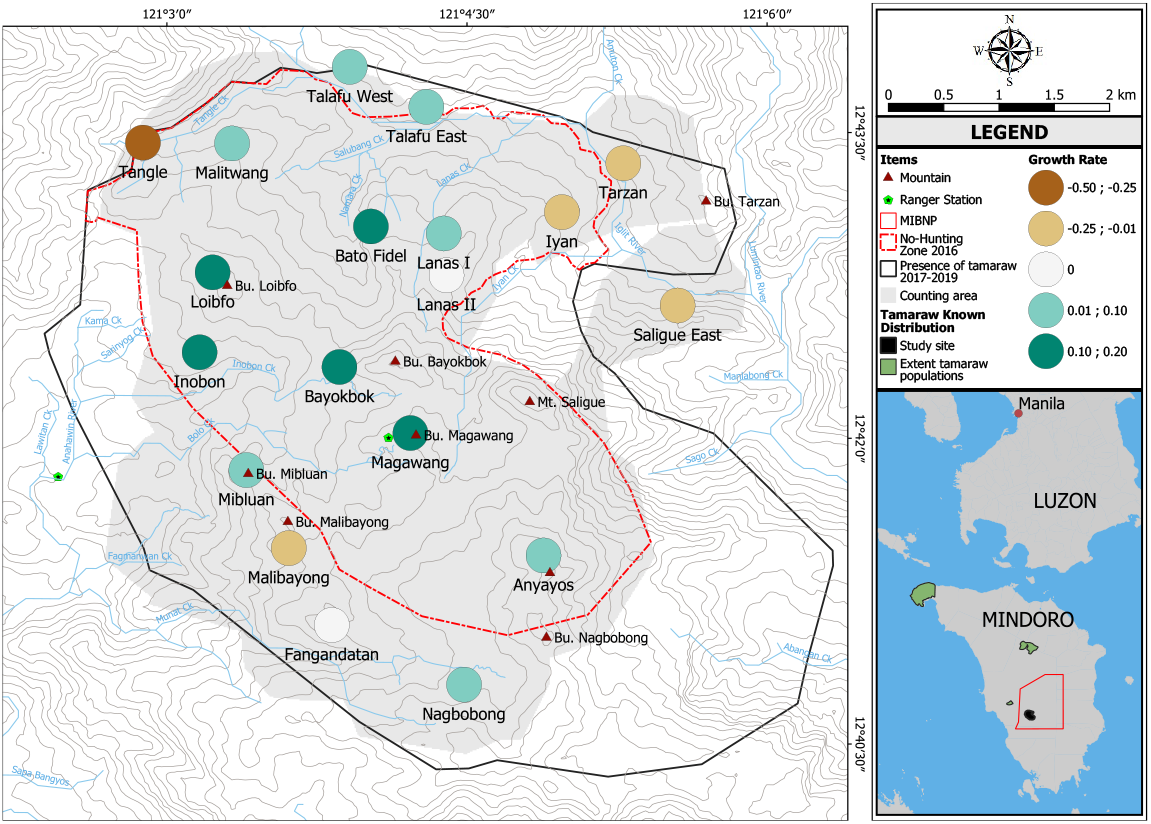
Spatial variation in the local growth rate of tamaraw (*Bubalus mindorensis*) abundance in Mounts Iglit-Baco Natural Park (MINBNP) on Mindoro island, Philippines, from years 2000 to 2021.

During the annual counts, individual tamaraws were counted simultaneously at the 18 vantage points for one hour and a half at dusk and dawn over a period of four consecutive days in late March or early April. For the 8 sessions and at each vantage point, at least two observers spotted and recorded all animals seen, writing down observation time, approximate location, and sex and age-category of individuals (split into calves, yearlings, sub-adults, and adults). This sampling design remained unchanged for the 21 years, though the number of observers could vary from year to year. Immediately after the completion of the 8 sessions, raw data were cross-checked and cleaned for potential multiple observations. Rangers observing at neighbouring vantage points decided together if a group of tamaraws was previously seen or not based on the timing, moving direction and composition of the group. When timing, moving direction and composition did match, then those individuals were withdrawn once from the total number of sightings. We derived the total number of different animals corrected for multiple counts for each vantage point and year, and constitute the consolidated counts C. These annual counts comprise the only available source of information about tamaraw population trends and dynamics for officials to use for conservation management decisions. Note that, because of the COVID-19 pandemic, the tamaraw counts in 2020 did not take place and we missed detailed counts (vantage point level) for years 2000 to 2003 and 2005 following the loss of field data sheets. We considered all these missing values as not available to be estimated by our statistical models (see below for details).

### Data analyses

For populations of long-lived and endangered species, robust and accurate estimations of the population growth rate (r) remain difficult to obtain. Knowledge about r requires either the estimation of demographic rates combined into a Leslie matrix (Leslie 1945) or the availability of long-term series of population abundance (Royama 1992), preferably on an annual basis. Because we were unable to mark tamaraw, both for legal and ethical reasons (Putman 1995), we could not individually identify animals, and thus it was not possible to apply methods of abundance estimations based on capture-recapture (see Schwarz & Seber 1999, for a review). We therefore used auto-regressive statistical models to analyze the annual counts. Working with abundance index data to derive an empirical estimates of r, however, comes with serious methodological issues (Îto 1972; Lindley 2003). Over the last decades, population counts have repeatedly been associated with a large sampling variance (*e.g*. Caughley 1977; Morellet *et al*. 2007), and coefficients of variation for population abundance close to 30% have been reported for various large mammal species (Lubow & Ransom 2016). From a conservation perspective, a large sampling variance in the input data for estimating *r* is a major pitfall because it leads to over-optimistic results, usually from an over-estimation of both the point estimate and its precision (Lindley 2003). Despite the availability of appropriate statistical methods to account for the sampling variance of population counts (Kalman filter, De Valpine & Hastings 2002), such tools remain seldom used in practice in conservation.

Here, we analyzed the consolidated number of observed tamaraws (*C*), as judged by observers to be different individuals, for each of the 18 vantage points, split by years (*C_s,t_*, where index *s* ∈ {1,…, 18} stands for the vantage point, and *t* ∈{2000,…, 2021}stands for the time in years. We implemented a state-space model to tease apart process from sampling variance, making the assumption that over- and under-counts cancel out and are randomly distributed from year to year. We worked in a Bayesian framework (e.g. Kéry & Schaub 2011) but note this is fully equivalent to a Kalman filter with a frequentist approach De Valpine & Hastings (2002). To do so, we defined *N_s,t_* as the unobserved ‘true’ abundance linked to *C_s,t_* by a random effect corresponding to the process variance 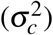 on the log scale to ensure positive values for abundance. Our baseline observation model was:

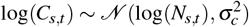

Accordingly, we computed the population growth rate *r_s,t_* of the tamaraw population at each vantage points *s*, as:

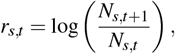

from counts corrected for the sampling variance, thus returning an unbiased estimation of the population growth rate provided there is no long-term trend in the detection probability of tamaraws. The overall population growth of the tamaraw population of MIBNP is then given by:

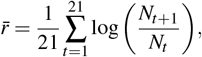

where 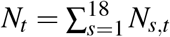. We input missing values at the vantage point level (years 2000–2003 and 2005) thanks to the sum constraint on *N_s,t_*, and the recursive equation linking *N_s,t_* and *N*_*s,t*+1_. By back-casting missing values of tamaraw abundance, we could estimate population growth at each vantage point (*r_s_*) with the same number of years and in turn, correctly test for the effect of human disturbance at the periphery of the CZM on the population dynamics.

To test our third hypothesis we investigated potential density-dependence in the annual growth rate by fitting two models classically used to detect negative density dependence (Dennis & Taper 1994): the Ricker and a Gompertz growth model, to be compared with an exponential growth model. The exponential growth model was fitted implicitly for the estimation of the average population growth rate 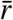 and served as our baseline model:

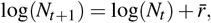

Then we fitted a Ricker growth model to the total number of tamaraws seen per year *N_t_* as follows:

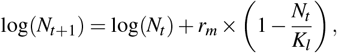

where *r_m_* is the maximum population growth for the tamaraw, and *K_l_* is the carrying capacity of the MIBNP. Likewise, we tested if the annual population growth rate would decrease in time with population density, or any other unmeasured variable varying in time, by fitting a particular formulation of the Gompertz model (Norton’s formulation, see Tjørve & Tjørve 2017):

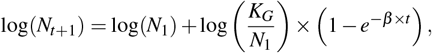

Because the pair of parameters *r_m_* and *K_l_* (Ricker) and *β* and *K_G_* (Gompertz) are not separately identifiable (Lebreton & Gimenez 2013), we used informative priors for *r_m_* and *β* based on the reported maximum population growth for other large bovids (see Traill *et al*. 2007, for a similar approach). Accordingly, we set a moderately informative prior value of mean 0.3 and a precision of 0.1 in our models (Table 1). We tested the occurrence of density-dependence processes in the MIBNP tamaraw population by computing a numerical approximation of the Bayes factor. We used MCMC samples to calculate the probability (*π_k_*) for our abundance time series to be generated by each of the three candidate demographic models during the model fit. At each MCMC step, a multinomial variable stated which of the exponential, Gompertz and Ricker model was the most likely underlying model given the data by taking the value one and zero otherwise. We estimated the three probabilities *π_k_* as the number of times each models received a one divided by the number of MCMC iterations.

**Table 1.** Estimated annual growth rates (r) of some large *bovinae* populations with body size comparable to the endangered tamaraw (*Bubalus mindorensis*). The observed population growth rate of tamaraws at Mounts Iglit-Baco Natural Park, Mindoro, Philippines, between 2000 and 2019 of *r* = 0.05 lies at the lower range of reported values. Note some populations may be in a very different ecological context (colonization, saturation) making comparisons challenging.

| Species | Location | *r* | Reference |
| --- | --- | --- | --- |
| *Bison bison* | Mackenzie Bison Sanctuary, Canada | 0.21 | Gates & Larter (1990) |
| *Bison bison* | Mackenzie Bison Sanctuary, Canada | 0.20 | Larter *et al.* (2000) |
| *Bison bison* | Yellowstone, Wyoming, USA | 0.07 | Fuller *et al.* (2007) |
| *Bison bison* | Yellowstone, Wyoming, USA | 0.30 | Singer & Norland (1994) |
| *Bison bonasus* | Białowieża, Poland | 0.07 | Myserud *et al.* (2007)<sup>1</sup> |
| *Bison bonasus* | Białowieża, Poland | 0.18 | Krasiński (1978)<sup>1</sup> |
| *Bos gaurus* | Thung Yai Naresuan Sanctuary, Thailand | 0.31 | Steinmetz *et al.* (2010) |
| *Bos javanicus* | Not applicable, model prediction | 0.32 | Hone *et al.* (2010) |
| *Bos javanicus* | Cobourg peninsula, Australia | 0.00 | Choquenot (1993) |
| *Boselaphus tragocamelus* | Kanha National Park, India | 0.18 | Mathur (1991) |
| *Boselaphus tragocamelus* | Keoladoe National Park, India | 0.02 - 0.06 | Haque (1990) |
| *Bubalus arnee* |  | 0.02 - 0.06 | Khatri *et al.* (2012) |
| *Bubalus arnee* | Koshi Tappu Wildlife Reserve, Nepal | 0.03 | Heinen & Paudel (2015) |
| *Bubalus bubalis* |  | 0.03 | Heinen (1993) |
| *Bubalus bubalis* |  | ? | Freeland and Boulton (1990) |
| *Bubalus mindorensis* | Mount Iglit-Baco National Park, Philippines | 0.05 | This study |
| *Pseudoryx nghetinhensis* | Vietnam & Laos | 0.05 | Kemp *et al.* (1997)<sup>b</sup> |
| *Syncerus caffer* | Hluhluwe-iMfolozi Park, South Africa | 0.12 | Jolles (2007) |
| *Syncerus caffer* | Zakouma National Park, Chad | 0.08 | Cornélis *et al.* (2014) |
| *Taurotragus oryx* | Lombard Nature Reserve, South Africa | 0.12 | Buys & Dott (1991)<sup>c</sup> |
| *Taurotragus oryx* | South Africa | 0.16 | Van Houtan *et al.* (2009) |
| *Taurotragus oryx* | Kruger National Park, South Africa | −0.02 | Nicholls *et al.* (1996) |
| *Taurotragus oryx* | Masai Mara Ecosystem, Kenya | −0.11 - 0.07 | Ottichilo *et al.* (2000) |
| *Taurotragus derbianus* | Senegal | 0.31 | Koláčková *et al.* (2011) |

We fitted our statistical model to the tamaraw count data with JAGS 4.0 (Plummer *et al*. 2003), with 3 MCMC chains and a burn-in of 40,000 iterations. We obtained parameter estimates with an additional 30 000 iterations and a thinning factor of 5 (therefore giving a total of 6 000 MCMC samples). We checked model convergence visually by investigating the mixing of MCMC chains, along with the 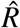 statistics that should be < 1.1 at convergence (Brooks & Gelman 1998). With the exception of *r_m_* (see above), we set uninformative priors with large variance for all other parameters to be estimated, and checked the sensitivity of our results to the average value of the priors by replicating the analyses with variable mean and initial values for prior distributions. We report all parameters estimates as the mean of the posterior distributions along with the 95% percentile for the credible intervals following Louis & Zeger (2008): _95%*low*_ *estimate* _95%*up*_.

Note that despite the fact that we analysed the number of tamaraws as described above, we chose not to display such numbers on figures and to report a standardized abundance 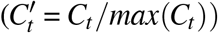 instead. This transformation applies to the estimated carrying capacities of the Gompertz and Ricker models, *K*′ = *K/max*(*C_t_*). We did not provide raw counts to avoid confusion between the index of relative tamaraw abundance we worked with, and a real population size estimator that was not implemented at MIBNP. With this variable transformation *K_l_* and *K_G_* are unitless and can no longer be interpreted as the maximum number of tamaraws but the maximum relative abundance 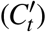 we can expect to observe at MIBNP. Reported values of the carrying capacities 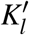 and 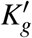 can hence take values >1 because the carrying capacity has not yet been reached. Biologically speaking, we put more emphasis on how far from the carrying capacity the current abundance of tamaraws is (*C_t_/K*) than of the number on animals *per se*.

## RESULTS

From our baseline model of tamaraw counts, we estimated an mean annual growth rate of 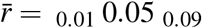 for the MIBNP population from 2000 to 2021, with an associated variance process equal to 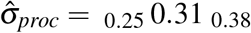 (Fig. 1). It means the observed abundance increased 2.41 times over the course of the study. As expected for count data, the sampling variance was large 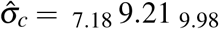, suggesting frequent double counts of the same individuals. For the vantage point specific growth rates (*r_s_*), we observed a marked variability in space (Fig. 2; see Supplementary Table S1) as shown by the fact that growth rates ranged between 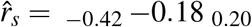 at Tarzan and 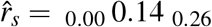 at Bato Fidel and Bayokbok vantage points. Without exception, 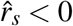 were located at the periphery of the no-hunting area (Iyan, Tarzan, Saligue East, Mibluan, Fangandatan), although 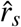 of a few vantage points were > 0 despite being located on the no-hunting zone border (Nagbobong, Malibayong, Talafu West and East). The mean growth rate outside of the no-hunting area was 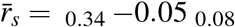 (9 vantage points) while tamaraw abundance increased at rate of 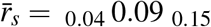 inside (9 vantage points).

We explicitly estimated the probabilities *π_k_* that the exponential, Ricker, or Gompertz models would be the underlying process generating the observed variation in tamaraw abundance (proxy of Bayes factor). The estimated probabilities for the three population dynamics models given our count data were 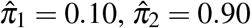 and 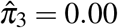, respectively, supporting our hypothesis of density-dependence feedbacks in the tamaraw population. Point estimates for the two parameters of the Ricker model were 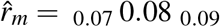 for the maximum population growth rate and 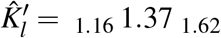 for the carrying capacity. Although the Gompertz model received no statistical support, the estimated coefficients were statistically significant and read 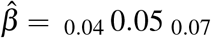, and 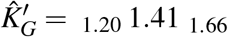 for the carrying capacity. Note that the statistical support for the Ricker model held for the dynamics of the core area only, that is, when considering only the 9 vantage points inside the no-hunting area (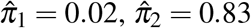 and 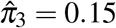).

## DISCUSSION

The isolated tamaraw population in MIBNP has been increasing in size since 2000 (Fig. 1). Undoubtedly, a positive growth rate of 0.05 over 21 years of monitoring is a noteworthy success for such a highly endangered species, both for local populations and for global conservation efforts. However, the devil lies in the details, and two additional results darken this otherwise positive picture. First, we found support for a source-sink dynamic of the tamaraw population linked to poaching and evidenced by a clear spatially structured pattern in the local population growth rates. Second, our results highlight a progressive decrease of the average growth rate in time, suggesting density-dependent effects at the population level. The recent population dynamics of tamaraw we document at Mindoro suggests that new conservation actions might be needed to secure the future of the species.

### Long-term Positive Growth of Tamaraws

The average population growth rate of 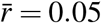 – corresponding to a 2.5 fold increase in abundance between 2000 and 2021 – gives strong evidence that successful conservation policies over the past two decades have led to an increase in tamaraw abundance. Nonetheless, an annual growth of 5% is relatively low compared to other similar-sized or even larger cattle species (Table 1). For instance, a population of African buffalo (*Syncerus caffer*) in South Africa grew at a rate of 12% per year for 28 years (Jolles 2007), and values close to 30% per year have regularly been recorded for bison (*Bison bison*) and banteng (Bos *javanicus*), despite the fact that these species are between 2 and 2.5 times larger in size than the tamaraw (Wilson & Mittermeier 2011). From this brief comparative approach, we would expect a substantially higher average growth rate for tamaraws if the population was in a colonizing phase. Given that the tamaraw is secure within the no-hunting area, our results suggest that specific environmental factors are limiting its growth in the CZM.

A first limiting factor could be related to the fact that the most available and accessible habitat for the tamaraw in MIBNP is grassland (83% of the CZM), which is dominated by competitive pioneer plant species such as cogon and wild sugarcane. Historical observations by Talbot & Talbot (1966) and more direct evidences (ES pers. obs.) at the *Gene Pool Farm* breeding center, Mindoro Island, also suggest that both invasive grasses make part of the tamaraw diet. Although these grasses likely provide tamaraws with abundant and high quality forage at the early stages of plant growth (Talbot & Talbot 1966), after burning for instance, in later stages of development these species are characterized by much lower nutritive value, especially during the dry season. Observations in the field indicate that once grass growth is complete, the stems exceeds tamaraw height (300cm vs. < 120 cm, respectively) and dry out, thus becoming both inaccessible and unpalatable. It is therefore possible that lack of food resource accessibility and availability could be limiting the growth rate for the population.

The annual burning of the grassland habitat for population count may also be impacting plant composition in the CZM, with implications for tamaraw population dynamics. Previous observations suggest that regular burning could promote the expansion of Siam weed (*Chromolaena odorata*, Nath *et al*. 2019), which appears to be rapidly spreading throughout our study site. The consumption of Siam weed by large herbivores is unclear. While some authors suggested potential toxicity of Siam weed leaves for consumers (Sajise *et al*.1974; Aterrado & Talatala-Sanico 1988), it makes a substantial part of the gaur *Bos gaurus* diet (Chaiyarat *et al*. 2021). Unlike its consumption, evidence for the avoidance of heavily invaded habitats by large herbivores is clearer. In African savannas, the induced changes in plant community, and the reduced detection of predators following the growth of Siam weed shifted habitat selection of grazing and browsing mammalian species (Rozen-Rechels *et al*. 2017). If, like other African large herbivores, tamaraw avoid Siam weed when foraging, its increasing dominance in grasslands at MIBNP could ultimately decrease the CZM carrying capacity. Although burning creates high-quality forage in the short-term, such an intensive habitat management practice is likely to cause a progressive decrease in habitat quality over the long term (see Dumalisile & Somers 2017, for an example of long-term effects of Siam weed on large herbivores in African savanna).

### Possible Impacts of Human Disturbance and Hunting

While the positive growth of the tamaraw population over the past two decades is evidence of a successful decrease in hunting pressure in the CZM, human activities could also account for the low tamaraw population growth rate we report. Tamaraw have always been traditionally hunted by indigenous people in Mindoro, and despite the “no-hunting agreement” established in 2016, both intentional and unintentional tamaraw deaths still occur. Moreover, the tamaraw is sought after for bushmeat by lowland poachers. Because illegal activities are difficult to control in MIBNP, it has likely led to increasing the mortality of all age-classes of tamaraws to an unknown, but perhaps considerable extent.

In the early years of the tamaraw monitoring, the apparent population growth (Fig. 1) likely resulted from immigrating individuals from outside of the CZM, and a high survival and reproduction rate inside. Today a different demographic process could be at work, with adult females gradually concentrating at the heart of the no-hunting area to avoid human disturbance. Conversely, a limited number of young individuals and adult males may venture outside of the no-hunting area, putting themselves at greater risk of dying. The net movements of animals in and out of the no-hunting area could lead to an apparent continuous growth of tamaraw abundance since the beginning of monitoring efforts, despite a shift in the underlying population dynamics.

The marked spatial heterogeneity we document, with negative growth rates at the periphery of the counting area, could indeed indicate a lower protection efficiency in areas further away from the main patrolling routes and ranger base camps. Such a result supports our second hypothesis, suggesting an unexpected contraction of the tamaraw range at MIBNP, and the disappearance of individuals at the periphery of the CZM. In other words, the CZM area may function like a source-sink system (Pulliam 1988), with the more central and better patrolled areas exhibiting positive growth rates, and more peripheral areas with higher hunting pressure serving as population sinks. Again, our findings provide some support for this idea, as local growth rates in the core area of the no-hunting zone have values as high as 14% (Fig. 2), which is more in line with the average growth rates that we reviewed for large bovids (Table 1).

Source-sink dynamics were found in populations of apex predators (wolves *Canis lupus:* O’Neil *et al*. (2020), lion *Panthera leo:* Mosser *et al*. (2009)) and other large herbivores of varying body size (red brocket *Mazama americana:*Naranjo & Bodmer (2007), *Giraffa camelopardalis:* Lee & Bolger (2017)) though at much larger spatial scales than for the tamaraw where source and sink areas are a couple of hundreds meters apart. The role of trophy or subsistence hunting (official or illegal) in source-sink dynamics is pervasive, particularly between protected and unprotected areas (Creel *et al*. 2016; Durant *et al*. 2017; Heurich *et al*. 2018), making human harvest the main target of conservation policies. Tamaraw is no exception and the observed source-sink dynamics stress the need for an improved control of poaching, and a further geographical expansion of the current 1 600ha no-hunting agreement area.

### Space Needed for Sustaining Population Growth

The gradual decrease of the tamaraw population growth rate in MIBNP, coupled with the overall abundance in the CZM, suggests a type of density-dependence is occurring, where a change in demographic rates lowers the number of individuals added to the population from year to year (Fowler 1987; Sinclair 1989). Predicted growth rates from the lowest to the highest abundance varied between *r =* 0.08 and *r =* 0.01 for tamaraw. From the current data, our best model suggests a carrying capacity of 1.4, meaning that only 40% of growth is left possible before reaching the carrying capacity. If competition for food or mating becomes strong then the main tamaraw population may experience an increased net loss of animals in the near future. Tamaraws may already be experiencing a food resource shortage, which could make the population more prone to parasitism (Stien *et al*. 2002) or epizootics (Davidson & Doster 1997). More generally, density-dependence should be of concerned and investigated for large herbivores living in protected environments lacking apex predators, which contribute to limit abundance and reduce annual growth of prey species (Ripple & Beschta 2012; Sinclair *et al*. 2003).

Another behaviour associated with high population abundance is dispersal (Matthysen 2005). In mammals, sub-adult individuals are more likely to disperse at high than low population abundance (e.g. Loe *et al*. 2009, for an example on red deer *Cervus elaphus*). At the high population abundance we documented in MIBNP, we would expect a greater number of dispersing individuals to venture outside of the no-hunting zone, where the risk of mortality is high because of hunting or poaching. In addition to jeopardizing the fate of the population by decreasing survival probabilities of sub-adults that are key to maintaining the future of the population, the empirical evidence for density-dependence that we report would limit the ability of this population to act as a source. With a combination of low recruitment and low survival of sub-adults, old individuals would quickly become over-represented in the population, which could depress growth even further.

Unfortunately, in the absence of individual monitoring of tamaraws to estimate demographic rates independently of abundance, the real cause of the density-dependence we report is uncertain. We are currently unable to determine if the gradual decrease in growth rate is the consequence of an increased mortality or dispersal rate, a reduced reproduction, or some combination of these factors. The difficulty of teasing apart different demographic processes is a well-known limit of the so-called pattern-oriented approach (*sensu* Krebs 2002) when working with time series of abundance. Further investigation of these possibilities is needed, as each situation would require different management approaches.

### Implications for Conservation

The main conservation goals for the tamaraw are (1) to ensure its long-term viability, and (2) support the expansion of the population within MIBNP and potentially in other sites. Current conservation measures assume that tamaraw population growth will continue in the next decade. The detection of density-dependence, however, suggests continued population growth will not be the case. In this situation, the dynamics of the tamaraw will change in the years to come if resources become the main limiting factor inside the no-hunting zone, forcing animals to disperse to, and potentially get killed in the outer zone. Several actions must be implemented by managers and authorities to address this issue in the short term. The priorities should be to expand the no-hunting zone or to create safety corridors through consensual agreement with indigenous people, both implemented in the context of creating a more sustainable land-use and traditional hunting system. Arriving at such an agreement with indigenous people will be a long process, but these conversations are already ongoing. Extending the no-hunting area will ultimately increase the sustainable population size, thus making it more viable and maintaining more genetic diversity. Likewise, allowing for dispersing individuals to leave the CZM and possibly settle in distant areas would relieve density dependence pressure on the core population.

However, one could argue that such conservation actions only represent a short-term solution, as even if the size of the no-hunting area was increased, the tamaraw population would eventually grow towards the carrying capacity. We acknowledge this, but believe that these short-term conservation actions are still important, as they would help buy valuable time that is necessary to devise more long-term strategies. Any long-term strategy will likely involve active population management through translocation. Two of the four known populations of tamaraw on Mindoro (Aruyan Malate and Mount Calavite) have extremely small population sizes, and are thus believed to be non-viable. Transporting a small number of animals to these populations would improve their viability, while simultaneously reducing the effects of density-dependence for the main population at MIBNP. The number of tamaraw that are translocated should be determined by the number of recruited individuals each year, so that ultimately there is no decrease in population size in the MIBNP. We acknowledge that translocation of tamaraw may be a perceived by stakeholders as a risky proposition, but a preliminary feasibility study for translocation and captive breeding is a realistic option and an appropriate next step. Ultimately, however, preventing the extinction of tamaraw will require decision makers to balance short-term actions with long-term strategies that involve active population management.

## Conclusion

Securing large areas of habitat is often an effective conservation approach, but may not always be a viable strategy to protect highly threatened species, as illustrated by our study on the tamaraw population in MIBNP. More generally, we show that positive growth rates could potentially hide a less optimistic picture for some threatened species. First, the growth of a population that is confined to a small conservation area may reflect a range contraction as a result of anthropogenic pressures, rather than the effect of increased abundance *per se*. Second, because of this, the population could face important demographic challenges that, in the long-term, lead to unusual problems for conservation. In this context, there is an increasing need to understand the interaction between anthropogenic activities and the population dynamics of threatened species. Our results suggest that, in some situations, conservation strategies designed to protect small, isolated populations may lead to long-term demographic issues that undermine species recovery. On the positive side, our findings also show how detailed analyses of relative population abundance data, even without absolute population size estimators, can help provide information that assists in the development of conservation measures that are essential for large mammal conservation.

## ACKNOWLEDGMENTS

We are grateful to Alvaro Gonzalez for his contribution to the fieldwork and his insights on tamaraw distribution at MIBNP. We also acknowledge the invaluable help of all rangers, students and volunteers who contributed to the annual tamaraw counts. Thanks are extended to 3 anonymous referees and to Jean-Michel Gaillard for their critical and constructive reading of the manuscript.

## SUPPLEMENTARY MATERIAL

**Table S1.** Estimates and 95% credible intervals of tamaraw local growth rate (r_s_) as observed on the 18 vantage points used to count the tamaraw population at Mounts Iglit-Baco National Park (Mindoro Island, Philippines) between 2000 and 2021.

| **Vantage point** | $r_s$ | **2.5%** | **97.5%** |
| --- | --- | --- | --- |
| Loibfo | 0.12 | −0.02 | 0.24 |
| Magawang | 0.11 | −0.02 | 0.24 |
| Bayokbok | 0.14 | 0.00 | 0.26 |
| Bato Fidel | 0.14 | 0.00 | 0.26 |
| Inubon | 0.11 | −0.02 | 0.24 |
| Mibluan | 0.08 | −0.06 | 0.22 |
| Nagbobong | 0.02 | −0.13 | 0.17 |
| Fangandatan | 0.00 | −0.19 | 0.16 |
| Anyayos | 0.07 | −0.07 | 0.20 |
| Lanas I | 0.07 | −0.08 | 0.21 |
| Iyan | −0.04 | −0.25 | 0.14 |
| Tarzan | −0.18 | −0.42 | 0.02 |
| Talafu East | 0.05 | −0.10 | 0.19 |
| Talafu West | 0.05 | −0.10 | 0.19 |
| Malitwang | 0.07 | −0.09 | 0.21 |
| Lanas II | 0.01 | −0.14 | 0.15 |
| Saligue East<sup>1</sup> | −0.08 | −0.40 | 0.15 |
| Malibayong | −0.01 | −0.20 | 0.15 |
| Tangle | −0.48 | −2.94 | 0.16 |

<sup>1</sup>Counts at this vantage point stopped in 2019, replaced by Tangle

## REFERENCES

Aterrado, E. & Talatala-Sanico, R. (1988). Status of *Chromolaena odorata* research in the Philippines. In: Proceedings of the first international Workshop on Biological Control of Chromolaena odorata, Bangkok, Thailand. pp. 53–55.

Beissinger, S.R. & Westphal, M.I. (1998). On the use of demographic models of population viability in endangered species management. The Journal of Wildlife Management, 62, 821–841.

Bonenfant, C., Gaillard, J.M., Coulson, T., Festa-Bianchet, M., Loison, A., Garel, M., Loe, L.E., Blanchard, P., Pettorelli, N., Owen-Smith, N., Du Toit, J. & Duncan, P. (2009). Empirical evidence of density-dependence in populations of large herbivores. Advances in Ecological Research, 41, 313–357.

Boyce, M.S. (1992). Population viability analysis. Annual Review of Ecology and Systematics, 23, 481–506.

Brooks, S.P. & Gelman, A. (1998). General methods for monitoring convergence of iterative simulations. Journal of Computational and Graphical Statistics, 7, 434–455.

Buys, D. & Dott, H. (1991). Population fluctuations and breeding of eland *Taurotragus oryx* in a western Transvaal nature reserve. Koedoe, 34, 31–36.

Caughley, G. (1977). Analysis of vertebrate populations. Wiley, London. Graeme Caughley. ‘A Wiley-Interscience Publication’ Includes index. Bibliography: p.217–228.

Caughley, G. (1994). Directions in conservation biology. Journal of Animal Ecology, 63, 215–244.

Chaiyarat, R., Prasopsin, S. & Bhumpakphan, N. (2021). Food and nutrition of gaur (bos gaurus ch smith, 1827) at the edge of khao yai national park, thailand. Scientific Reports, 11, 1–11.

Choquenot, D. (1991). Density-dependent growth, body condition, and demography in feral donkeys: testing the food hypothesis. Ecology, 72, 805–813.

Choquenot, D. (1993). Growth, body condition and demography of wild banteng (Bos *javanicus*) on Cobourg Peninsula, northern Australia. Journal of Zoology, 231, 533–542.

Cornélis, D., Melletti, M., Korte, L., Ryan, S.J., Mirabile, M., Prin, T. & Prins, H.H. (2014). African buffalo Syncerus caffer (Sparrman, 1779), Cambridge University Press, pp. 326–372.

Courchamp, F., Clutton-Brock, T.H. & Grenfell, B. (1999). Inverse density dependence and the Allee effect. Trends in Ecology & Evolution, 14, 405–410.

Creel, S., M’soka, J., Dröge, E., Rosenblatt, E., Becker, M.S., Matandiko, W. & Simpamba, T. (2016). Assessing the sustainability of african lion trophy hunting, with recommendations for policy. Ecological Applications, 26, 2347–2357.

Custodio, C.C., Lepiten, M.V. & Heaney, L.R. (1996). Bubalus mindorensis. Mammalian Species, pp. 1–5.

Davidson, W. & Doster, G. (1997). Health characteristics and white-tailed deer population density in the southeastern United States. In: The Science of overabundance: deer ecology and population management (eds. McShea, W.J., Underwood, H.B. & Rappole, J.H.). Smithsonian Institution Press, Washington, DC (USA), pp. 164–184.

De Valpine, P. & Hastings, A. (2002). Fitting population models incorporating process noise and observation error. Ecological Monographs, 72, 57–76.

Dennis, B. & Taper, M. (1994). Density dependence in time series observations of natural populations: estimation and testing. Ecological Monographs, 64, 205–224.

Dumalisile, L. & Somers, M.J. (2017). The effects of an invasive alien plant (*Chromolaena odorata*) on large African mammals. Nature Conservation Research, 2, 102–108.

Durant, S.M., Mitchell, N., Groom, R., Pettorelli, N., Ipavec, A., Jacobson, A.P., Woodroffe, R., Böhm, M., Hunter, L.T., Becker, M.S. et al. (2017). The global decline of cheetah acinonyx jubatus and what it means for conservation. Proceedings of the National Academy of Sciences, 114, 528–533.

Eberhardt, L.L. (1977). Optimal policies for conservation of large mammals, with special references to marine ecosystems. Environmental Conservation, 4, 205–212.

Eberhardt, L.L. (2002). A paradigm for population analysis of long-lived vertebrates. Ecology, 83, 2841–2854.

Fowler, C.W. (1987). A review of density dependence in populations of large mammals. Current Mammalogy, 1, 401–441.

Fuller, J.A., Garrott, R.A., White, P., Aune, K.E., Roffe, T.J. & Rhyan, J.C. (2007). Reproduction and survival of Yellowstone bison. Journal of Wildlife Management, 71, 2365–2372.

Gates, C. & Larter, N. (1990). Growth and dispersal of an erupting large herbivore population in northern Canada: the Mackenzie wood bison (*Bison bison athabascae*). Arctic, pp. 231–238.

Haque, N. (1990). Study on the ecology of wild ungulates of Keoladeo national Park Bharatpur, Rajasthan. Ph.D. thesis, Aligarh Muslim University.

Heinen, J.T. & Paudel, P.K. (2015). On the translocation of wild asian buffalo *Bubalis arnee* in Nepal: are feral backcrosses worth conserving? Conservation Science, 3, 11–19.

Henle, K., Sarre, S. & Wiegand, K. (2004). The role of density regulation in extinction processes and population viability analysis. Biodiversity and Conservation, 13, 9–52.

Heude, P. (1888). Éludes sur les ruminants de l’Asie orientale. cerfs des Philippines et de l’Indo-Chine. Mémoires Concernant l’Histoire Naturelle de l’Empire Chinois par des Pères de la Compagnie de Jésus, 2, 1–10.

Heurich, M., Schultze-Naumburg, J., Piacenza, N., Magg, N., Červenỳ, J., Engleder, T., Herdtfelder, M., Sladova, M. & Kramer-Schadt, S. (2018). Illegal hunting as a major driver of the source-sink dynamics of a reintroduced lynx population in central europe. Biological conservation, 224, 355–365.

Hone, J. & Clutton-Brock, T.H. (2007). Climate, food, density and wildlife population growth rate. Journal of Animal Ecology, 76, 367–367.

Hone, J., Duncan, R.P. & Forsyth, D.M. (2010). Estimates of maximum annual population growth rates (*rm*) of mammals and their application in wildlife management. Journal of Applied Ecology, 47, 507–514.

Îto (1972). On the methods for determining density-dependence by means of regression. Oecologia, 10, 776–778.

Jolles, A.E. (2007). Population biology of african buffalo (*Syncerus caffer*) at Hluhluwe-iMfolozi Park, South Africa. African Journal of Ecology, 45, 398.

Kemp, N., Dilger, M., Burgess, N. & Van Dung, C. (1997). The saola *Pseudoryx nghetinhensis* in Vietnam – new information on distribution and habitat preferences, and conservation needs. Oryx, 31, 37–44.

Kéry, M. & Schaub, M. (2011). Bayesian population analysis using WinBUGS: a hierarchical perspective. Academic Press, Amsterdam, the Netherlands.

Koláčková, K., Hejcmanova, P., Antonínová, M. & Brandl, P. (2011). Population management as a tool in the recovery of the critically endangered Western Derby eland *Taurotragus derbianus* in Senegal, Africa. Wildlife Biology, 17, 299–311.

Krasiński, Z.A. (1978). Dynamics and structure of the european bison population in the białowieza Primeval Forest. Acta Theriologica, 23, 3–48.

Krebs, C.J. (2002). Two complementary paradigms for analysing populations dynamics. Philosophical Transactions of the Royal Society of London, Series B, 357, 1211–1219.

Krüger, O. & Lindström, J. (2001). Habitat heterogeneity affects population growth in goshawk *Accipiter gentilis*. Journal of Animal Ecology, 70, 173–181.

Lande, R. (1993). Risks of population extinction from demographic and environmental stochasticity and random catastrophes. American Naturalist, 142, 911–927.

Langwig, K.E., Frick, W.F., Bried, J.T., Hicks, A.C., Kunz, T.H. & Marm Kilpatrick, A. (2012). Sociality, density-dependence and microclimates determine the persistence of populations suffering from a novel fungal disease, white-nose syndrome. Ecology Letters, 15, 1050–1057.

Larter, N., Sinclair, A., Ellsworth, T., Nishi, J. & Gates, C. (2000). Dynamics of reintroduction in an indigenous large ungulate: the wood bison of northern Canada. Animal Conservation, 3, 299–309.

Lebreton, J.D. & Gimenez, O. (2013). Detecting and estimating density dependence in wildlife populations. Journal of Wildlife Management, 77, 12–23.

Lee, D.E. & Bolger, D.T. (2017). Movements and source–sink dynamics of a masai giraffe metapopulation. Population Ecology, 59, 157–168.

Leslie, P.H. (1945). On the use of matrices in population mathematics. Biometrika, 33, 182–212.

Lindley, S.T. (2003). Estimation of population growth and extinction parameters from noisy data. Ecological Applications, 13, 806–813.

Loe, L.E., Mysterud, A., Veiberg, V. & Langvatn, R. (2009). Negative density-dependent emigration of males in an increasing red deer population. Proceedings of the Royal Society B: Biological Sciences, 276, 2581–2587.

Long, B., Schütz, E., Appleton, M., Gonzales, A., Holland, J., Lees, C., Marandola, E., Natural, V.C.J., Pineda-David, M.T.J., Salao, C., Slade, J., Tan, E.H.P., Tiongson, L. & Young, S. (2018). Review of tamaraw (*Bubalus mindorensis*) status and conservation actions. BULLetin, 1, 18–34.

Louis, T.A. & Zeger, S.L. (2008). Effective communication of standard errors and confidence intervals. Biostatistics, 10, 1–2.

Lubow, B.C. & Ransom, J.I. (2016). Practical bias correction in aerial surveys of large mammals: Validation of hybrid double-observer with sightability method against known abundance of feral horse (*Equus caballus*) populations. PLoS One, 11, e0154902.

Maala, C.P. (2001). Endangered Philippine wildlife species with special reference to the Philippine eagle (*Pithecophaga jefferyi*) and tamaraw (*Bubalus mindorensis*). Journal of International Development and Cooperation, 8, 1–17.

Mace, G.M., Collar, N.J., Gaston, K.J., Hilton-Taylor, C., Akçakaya, H.R., Leader-Williams, N., Milner-Gulland, E.J. & Stuart, S.N. (2008). Quantification of extinction risk: IUCN’s system for classifying threatened species. Conservation Biology, 22, 1424–1442.

Mathur, V.B. (1991). The ecological interaction between habitat composition, habitat quality and abundance of some wild ungulates in India. Ph.D. thesis, University of Oxford.

Matsubayashi, H., Boyles, R.M., Salac, R.L., Del Barrio, A., Cruz, L., Garcia, R.A., Ishihara, S. & Kanai, Y. (2010). Present status of tamaraw (*Bubalus mindorensis*) in Mt. Aruyan, Mindoro, Philippines. Tropics, 18, 167–170.

Matthysen, E. (2005). Density-dependent dispersal in birds and mammals. Ecography, 28, 403–416.

Morellet, N., Gaillard, J.M., Hewison, A.J.M., Ballon, P., Boscardin, Y., Duncan, P., Klein, F. & Maillard, D. (2007). Indicators of ecological change: new tools for managing populations of large herbivores. Journal of Applied Ecology, 44, 634–643.

Mosser, A., Fryxell, J.M., Eberly, L. & Packer, C. (2009). Serengeti real estate: density vs. fitness-based indicators of lion habitat quality. Ecology Letters, 12, 1050–1060.

Mugabo, M., Perret, S., Legendre, S. & Le Galliard, J.F. (2013). Density-dependent life history and the dynamics of small populations. Journal of Animal Ecology, 82, 1227–1239.

Mysterud, A., Bartoń, K.A., Jedrzejewska, B., Krasiński, Z.A., Niedziałkowska, M., Kamler, J.F., Yoccoz, N.G. & Stenseth, N.C. (2007). Population ecology and conservation of endangered megafauna: the case of european bison in Białowieża Primeval Forest, Poland. Animal Conservation, 10, 77–87.

Naranjo, E.J. & Bodmer, R.E. (2007). Source–sink systems and conservation of hunted ungulates in the lacandon forest, mexico. Biological Conservation, 138, 412–420.

Nath, A., Sinha, A., Lahkar, B.P. & Brahma, N. (2019). In search of aliens: Factors influencing the distribution of *Chromolaena odorata* L. and Mikania micrantha Kunth in the Terai grasslands of Manas National Park, India. Ecological Engineering, 131, 16–26.

Neel, M.C., Leidner, A.K., Haines, A., Goble, D.D. & Scott, J.M. (2012). By the numbers: How is recovery defined by the US endangered species act? BioScience, 62, 646–657.

Nicholls, A., Viljoen, P., Knight, M. & Van Jaarsveld, A. (1996). Evaluating population persistence of censused and unmanaged herbivore populations from the Kruger National Park, South Africa. Biological Conservation, 76, 57–67.

Nicholson, A.J. (1933). The balance of animal populations. Journal of Animal Ecology, S2, 132–178.

O’Neil, S.T., Vucetich, J.A., Beyer Jr, D.E., Hoy, S.R. & Bump, J.K. (2020). Territoriality drives preemptive habitat selection in recovering wolves: Implications for carnivore conservation. Journal of Animal Ecology, 89, 1433–1447.

Ottichilo, W.K., De Leeuw, J., Skidmore, A.K., Prins, H.H. & Said, M.Y. (2000). Population trends of large non-migratory wild herbivores and livestock in the Masai Mara ecosystem, Kenya, between 1977 and 1997. African Journal of Ecology, 38, 202–216.

Pancho, J. & Plucknett, D. (1971). *Chromolaena odorata* (l.) rm king and h. robinson – a new record of a noxious weed in the Philippines. Philippine Journal of Animal Science, 8, 143–149.

Peterson, C.H. & Black, R. (1988). Density-dependent mortality caused by physical stress interacting with biotic history. American Naturalist, 131, 257–270.

Plummer, M. et al. (2003). JAGS: A program for analysis of bayesian graphical models using gibbs sampling. In: Proceedings of the 3rd international workshop on distributed statistical computing. Vienna, Austria, vol. 124, p. 10.

Pulliam, H.R. (1988). Sources, sinks, and population regulation. The American Naturalist, 132, 652–661.

Putman, R. (1995). Ethical considerations and animal welfare in ecological field studies. Biodiversity & Conservation, 4, 903–915.

Ripple, W.J. & Beschta, R.L. (2012). Large predators limit herbivore densities in northern forest ecosystems. European Journal of Wildlife Research, 58, 733–742.

Royama, T. (1992). Analytical population dynamics. Chapman & Hall.

Rozen-Rechels, D., te Beest, M., Dew, L.A., le Roux, E., Druce, D.J. & Cromsigt, J.P. (2017). Contrasting impacts of an alien invasive shrub on mammalian savanna herbivores revealed on a landscape scale. Diversity and Distributions, 23, 656–666.

Sajise, P., Palis, R., Norcio, N., Lales, J. et al. (1974). The biology of *Chromolaena odorata* (L.) RM King and H. Robinson. 1. flowering behaviour, pattern of growth and nitrate metabolism. Philippine Weed Science Bulletin, 1, 17–24.

Schwarz, C. & Seber, G.A.F. (1999). A review of estimating animal abundance III. Statistical Science, 14, 427–456.

Shaffer, M.L. (1981). Minimum population sizes for species conservation. BioScience, 31, 131–134.

Sinclair, A.R., Mduma, S. & Brashares, J.S. (2003). Patterns of predation in a diverse predator-prey system. Nature, 425, 288–290.

Sinclair, A.R.E. (1989). Population regulation in animals. In: Ecological concepts: the contribution of ecology to an understanding of the natural world (eds. Cherrett, J.M. & Bradshaw, A.D.). Blackwell Scientific Publications, Oxford, England., pp. 197–241.

Singer, F.J. & Norland, J.E. (1994). Niche relationships within a guild of ungulate species in Yellowstone National Park, Wyoming, following release from artificial controls. Canadian Journal of Zoology, 72, 1383–1394.

Steinmetz, R., Chutipong, W., Seuaturien, N., Chirngsaard, E. & Khaengkhetkarn, M. (2010). Population recovery patterns of southeast asian ungulates after poaching. Biological Conservation, 143, 42–51.

Stephens, P.A., Sutherland, W.J. & Freckleton, R.P. (1999). What is the Allee effect? Oikos, pp. 185–190.

Stien, A., Irvine, R.J., Ropstad, E., Halvorsen, O., Langvatn, R. & Albon, S.D. (2002). The impact of gastrointestinal nematodes on wild reindeer: experimental and cross-sectional studies. Journal of Animal Ecology, 71, 937–945.

Talbot, L.M. & Talbot, M.H. (1966). The tamarau (*Bubalus mindorensis* (heude)). observations and recommendations. Mammalia, 30, 1–12.

Tjørve, K.M. & Tjørve, E. (2017). The use of Gompertz models in growth analyses, and new Gompertz-model approach: An addition to the unified-Richards family. PloS one, 12, e0178691.

Traill, L.W., Bradshaw, C.J.A. & Brook, B.W. (2007). Minimum viable population size: a meta-analysis of 30 years of published estimates. Biological Conservation, 139, 159–166.

Van Houtan, K.S., Halley, J.M., Van Aarde, R. & Pimm, S.L. (2009). Achieving success with small, translocated mammal populations. Conservation Letters, 2, 254–262.

Van der Wal, E., Laforge, M.P. & McLoughlin, P.D. (2014). Density dependence in social behaviour: home range overlap and density interacts to affect conspecific encounter rates in a gregarious ungulate. Behavioral Ecology and Sociobiology, 68, 383–390.

Wilson, D. & Mittermeier, R. (2011). Handbook of the mammals of the world. Vol. 2. Hoofed Mammals. Lynx Edicions, Barcelona.

